# Polo-like kinase 1 regulates immune synapse assembly and cytotoxic T cell function by driving microtubule dynamics

**DOI:** 10.1101/2023.07.12.548674

**Authors:** Fabrizia Zevolini, Anna Onnis, Roxana Khazen, Sabina Müller, Giuseppe Marotta, Salvatore Valitutti, Francesca Finetti, Cosima T Baldari

## Abstract

Elimination of virally infected or tumoral cells is mediated by cytotoxic T cells (CTL). Upon antigen recognition CTLs assemble a specialized signaling and secretory domain at the interface with their target, the immune synapse (IS). During IS formation CTLs acquire a transient polarity, marked by re-orientation of the centrosome and microtubule cytoskeleton toward the IS, thus directing the transport and delivery of the lytic granules to the target cell. Based on the implication of the kinase Aurora-A in CTL function we hypothesized that its substrate, the mitotic regulator Polo-like kinase 1 (PLK1), may participate in CTL IS assembly. We demonstrate that PLK1 is phosphorylated upon TCR triggering and polarizes to the IS. PLK1 silencing or inhibition results in impaired IS assembly and function, as witnessed by defective synaptic accumulation of TCRs as well as compromised centrosome and lytic granule polarization to the IS, resulting in impaired target cell killing. This function is achieved by coupling early signaling to microtubule dynamics, a function pivotal for CTL-mediated cytotoxicity. These results identify PLK1 as a new player in CTL IS assembly and function.

**Summary statement:** The mitotic kinase Polo-like kinase 1 promotes centrosome polarization to the immune synapse in cytotoxic T cells by coupling TCR signaling to microtubule dynamics

## Introduction

Cytotoxic T cells (CTL) are the arm of adaptive immunity specialized in the elimination of virally infected or tumoral cells. To achieve this function CTLs have evolved three complementary mechanisms that function sequentially to allow for the serial killing of cognate targets. The first involves the lytic granules (LG), which are specialized secretory lysosomes enriched in the pore-forming toxin perforin (Prf) and the proteolytic enzymes granzymes (Gzm) packed on a serglycin scaffold to form a dense core that is solubilized upon LG exocytosis. Gzms are then delivered to cell targets through pores formed by Prf polymerization to orchestrate their apoptotic demise (Cassioli and Baldari, 2022; McKenzie and Valitutti, 2023). The second, which is essential for the sustained killing activity of CTLs when their LG stores become exhausted, is mediated by the death receptor ligand FasL, that triggers the apoptotic cascade initiated by caspase 8 through interaction with Fas on target cells (Green and Llambi, 2015; Cassioli and Baldari, 2022). A third mechanism that has been recently discovered involves the release of supramolecular attack particles (SMAP), autonomous killing entities that are composed of a core contaning Prf and Gzms enclosed in a non-membranous glycoprotein shell (Balint et al., 2020) and stored in a different type of LG, the multicore cytotoxic granule (Chang et al., 2022; Cassioli and Baldari, 2022).

CTL-mediated killing requires the assembly of the immune synapse (IS), a specialized signaling and secretory platform that forms at the cell target interface (De La Roche et al., 2016). TCR interaction with MHC-bound cognate peptide antigen on the target cell surface triggers a signaling cascade that promotes the translocation of the centrosome and associated microtubule cytoskeleton beneath the synaptic membrane, resulting in the acquisition by the CTL of a transient polarity (Gomez and Billadeau, 2008; Huse et al., 2013). This polarity is essential for the directional transport of the LGs and FasL-enriched vesicles towards the target cell and their precise delivery at the IS, thus ensuring the selective killing of the cognate target.

A tight interplay of the tubulin and actin cytoskeletons coordinates the movement of the centrosome towards the IS, with microtubule (+) ends interacting with F-actin at the IS periphery with the assistance the adaptors IQ-GAP, ezrin and Cdc42IP (Banerjee et al., 2007; Gorman et al., 2012; Ilani et al., 2007). Regulators of microtubule and actin dynamics, including the dynein/dynactin motor, the microtubule end-binding protein EB1, the small GTPase Rac1, the formins mDia and INF2, and the histone deacetylase HDAC6 also contribute to this process (Andrés-Delgado et al., 2013; Bouchet et al., 2016; Gomez and Billadeau, 2008; Martín-Cófreces et al., 2012; Sanchez et al., 2019; Serrador et al., 2004). Centrosome positioning at the IS center is paralleled by F-actin clearance to leave an actin-poor region that facilitates LG exocytosis (Ritter et al., 2015). The signals that regulate centrosome polarization in response to TCR triggering are as yet only partly defined. PLCγ activation downstream of early TCR signaling has been identified as a key initiating event. Diacyl glycerol produced by PLCγ at the IS center acts as a polarity determinant to attract the atypical PKCs such as ζ (Huse et al., 2013). These kinases orchestrate the process of centrosome translocation through phosphorylation of regulators of cytoskeleton dynamics such as myosin II, that interacts with the dynein/dynactin complex to provide the pulling forces required for centrosome movement (Liu et al., 2013).

Interestingly, the mitotic serine-threonine kinase Aurora A (AurA) has been recently implicated in the signaling events leading to IS formation in CD4^+^ T cells (Blas-Rus et al., 2016). In mitotic cells AurA expression and activity peak in late G2, when the protein is concentrated at centrosomes (Bischoff, 1998). During mitosis AurA regulates centrosome and spindle dynamics by recruiting microtubule nucleation and stabilization factors (Sardon et al., 2008). In non-dividing CD4^+^ T cells AurA is activated upon TCR stimulation and modulates TCR signaling to regulate the growth of the microtubules arising from the centrosome, thereby facilitating the polarized vesicular traffic that is essential for T cell activation and effector function (Blas-Rus et al., 2016). Here we have investigated the role of the mitotic kinase PLK1, a key substrate of AurA controlling centrosome and microtubule dynamics during cell cycle progression (Joukov and De Nicolo, 2018), in CTL IS assembly. We show that PLK1 is associated with the centrosome and is co-mobilized to the IS, where a pool of active PLK1 associates with the synaptic membrane. We provide evidence that PLK1 subserves AurA-dependent and -independent functions in CTLs, where it promotes centrosome polarization to the IS and the downstream events leading to target cell killing by coupling TCR signaling to microtubule growth. Our results identify PLK1 as a new player in CTL IS assembly and effector function.

## Results

### PLK1 is recruited to the CTL IS together with the centrosome and is activated in response to TCR signaling

PLK1 has been reported to localize at the centrosome in a variety of cell types (Colicino and Hehnly, 2018), hyet its localization in T lymphocytes is presentòy elusive. To address this point we generated CTLs from peripheral blood CD8^+^ T cells freshly purified from healthy donors and activated using beads coated with anti-CD3 and anti-CD28 mAbs in the presence of IL-2. Under these conditions CD8^+^ T cells differentiate to fully functional CTLs by day 5-7 (Onnis et al., 2023).

Immunoblot analysis showed that PLK1 is expressed in freshly purified CD8^+^ T cells and that its expression is increased during their differentiation to CTLs (Fig.1A). To determine the intracellular localization of PLK1 in CTLs we carried out a confocal microscopy analysis of CTLs co-stained for PLK1 and either pericentriolar material 1 (PCM1), a centriolar satellite protein that is responsible for PLK1 recruitment to the pericentriolar matrix (Wang et al., 2013), or the centrosome marker γ-tubulin. Similar to other cell types, PLK1 was found to localize at the centrosome in CTLs (Fig.1B). PLK1 also co-localized with the centrosome in CMV-specific T cell clones (Fig.S1A,B) and in other T cell settings, including Jurkat T cells and freshly purified peripheral blood CD4^+^ T cells (Fig.S2A). AurA also displayed a centrosomal localization in CTLs (Fig.S3A,B), as previously reported for CD4^+^ T cells (Blas-Rus et al., 2016).

**Figure 1.**
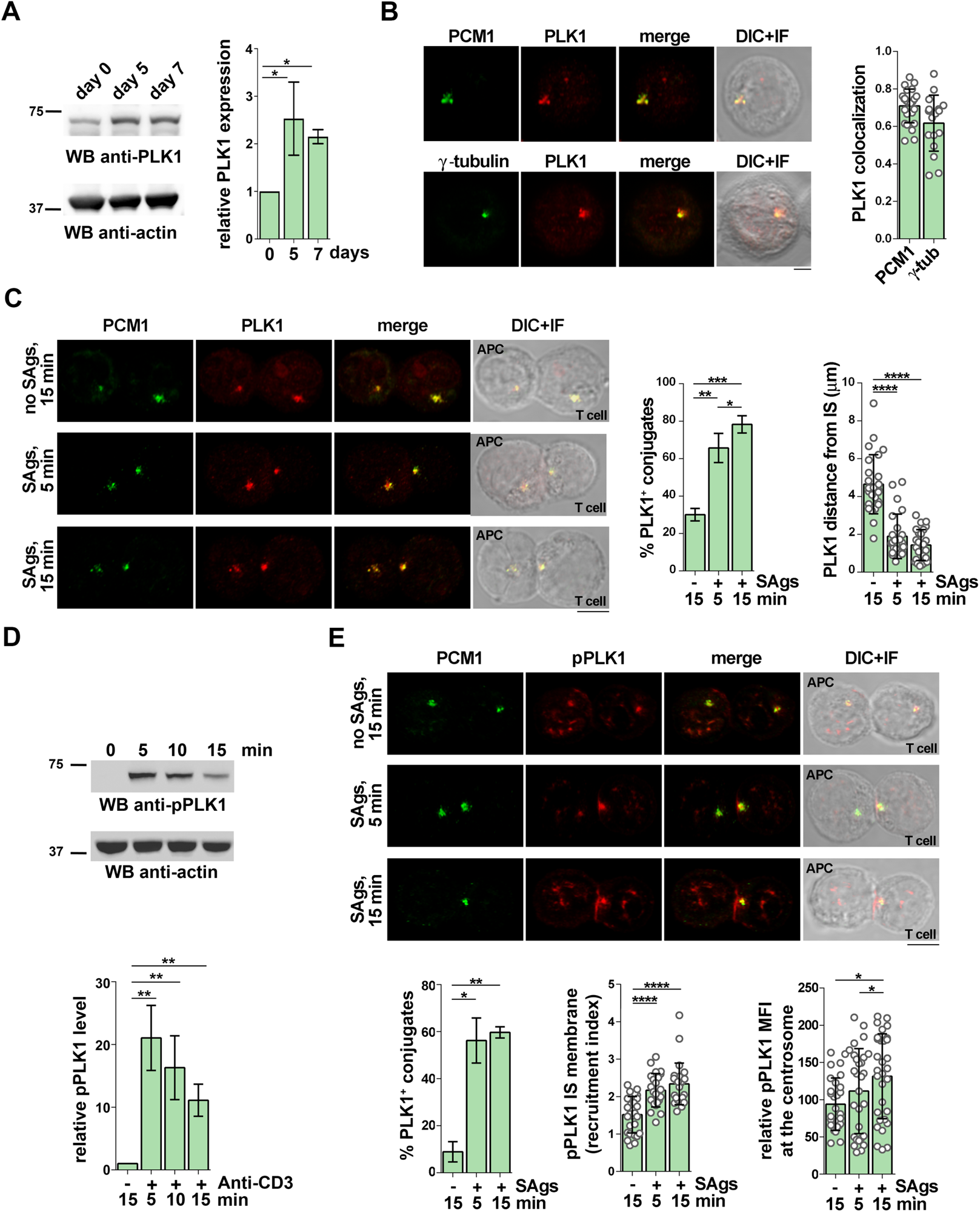
PLK1 is recruited to and phosphorylated at the IS in CTLs. **(A)** Immunoblot analysis with anti-PLK1 antibody of lysates of CD8^+^ T cells collected at days 0, 5 or 7 following stimulation with anti-CD3/CD28 coated beads. A control anti-actin blot of the stripped filter is shown below. The migration of molecular mass markers is indicated. The quantification of PLK1 in CTL lysates is reported in the graph (n=4, mean fold ± SD; ANOVA). **(B)** Quantification using Mander’s coefficient of the weighted colocalization of PLK1 and the centrosome markers PCM1 or γ-tubulin in CTLs (30 cells/sample, n=3; mean fold ± SD). Medial optical sections of representative images are shown. Scale bar: 5 μm. **(C)** Immunofluorescence analysis of PLK1 and PCM1 5 or 15 minutes after conjugate formation of CTLs and SAg-loaded Raji cells. Conjugates formed in the absence of SAgs (no SAgs) were used as negative control. Scale bar: 5 μm. The graphs show the percentage of conjugates with PLK1 polarization at the IS (15 cells/sample, n=3; ANOVA)(left) and the quantification of the distance of PLK1 from the IS membrane (μm)(20 cells/sample, n=3; Kurskall Wallis test)(right), respectively. **(D)** Immunoblot analysis with anti-pPLK1 antibody of lysates of CTLs either unstimulated or stimulated with anti-CD3 mAb for the indicated times. A control anti-actin blot of the stripped filter is shown below. The migration of molecular mass markers is indicated. The quantification of pPLK1 in anti-CD3 mAb-stimulated or unstimulated CTLs is reported in the graph (n=3, mean fold ± SD; one sample t test). **(E)** Immunofluorescence analysis of p-PLK1 and PCM1 5 or 15 minutes after conjugate formation of CTLs and SAg-pulsed Raji B cells. Scale bar: 5 μm. The graphs show the percentage of conjugates harbouring accumulation of the phosphorylated form of PLK1 at the IS region (15 cells/sample, n=3; mean ± SD, ANOVA)(left), the quantification of pPLK1 intensity at the IS membrane *versus* the total cell membrane (center) and the quantification of pPLK1 fluorescence intensity in the centrosome region defined from the point of PCM1 maximal intensity (24 cells/sample, n=3; mean ± SD, Kurskall Wallis test)(right).*P < 0.05; **P < 0.01; ***P < 0.001; ****P < 0.0001.

To assess whether PLK1 could participate in centrosome dynamics during IS formation, we carried out a similar co-localization analysis in conjugates of CTLs with Raji B cells pulsed with a mix of staphylococcal superantigens (SAgs) allowing for the polyclonal activation of T cells independently of antigen specificity. The analysis was carried out either 5 or 15 minutes after conjugate formation to visualize the events taking place during IS assembly (De La Roche et al., 2016). PLK1, as well as AurA, were found to localize at the CTL subsynaptic region in association with the centrosome as early as 5 min after conjugate formation (Fig.1C; Fig.S3A,C). Similar results were obtained in antigen-specific conjugates formed using CMV-specific CD8^+^ T cell clones (Fig.S1A,C). PLK1 co-polarized with the centrosome to the IS also in Jurkat T cells and freshly purified peripheral blood CD4^+^ T cells (Fig.S2B,C).

The kinase activity of PLK1 depends on its phosphorylation by AurA and its co-factor Bora on the T-loop Thr210 residue (Bruinsma et al., 2013). Immunoblot analysis of the phosphorylated, active form of PLK1 (pPLK1) revealed that, while pPLK1 was barely detectable in unstimulated cells, there was a sharp increase in response to TCR engagement, with the highest levels detected at the earliest time point analyzed (Fig.1D). To assess the specific localization of activated PLK1 at the IS, we performed a confocal microscopy analysis of pPLK1 in CTLs conjugated with either SAg-pulsed or unpulsed Raji B cells. pPLK1 could be detected at the centrosome in conjugates formed both in the absence of and in the presence of SAgs (Fig.1E). Additionally, pPLK1 was detectable at the IS in SAg-specific conjugates (Fig.1E). Hence PLK1 polarizes to the IS in association with the centrosome and a pool of its active form accumulates at the synaptic membrane, suggesting a potential implication in CTL IS formation.

### PLK1 is required for IS formation and TCR signaling in CTLs

IS formation is marked by the accumulation of TCR complexes at the central area of the T cell interface with the APC. Two TCR pools, of which one associated with the plasma membrane and the other with recycling endosomes, are sequentially recruited to the IS following TCR engagement by cognate antigen, thereby ensuring sustained TCR signaling (Soares et al., 2013). To investigate the potential role of PLK1 at the CTL IS we took advantage of the highly selective ATP analogue BI2536, which inhibits PLK1 activity at low nanomolar concentrations (Steegmaier et al., 2007). Confocal microscopy analysis of SAg-specific CTL conjugates revealed that the synaptic accumulation of TCRs was impaired in BI2536-treated CTLs, as assessed by labelling the CD3ζ chain of the TCR complex (Fig.2A), indicating that PLK1 is required for IS formation in CTLs. Similar results were obtained on Jurkat and freshly purified peripheral CD4^+^ T cells (Fig.S2D,E). To rule out off-target effects of BI2536, TCR accumulation at the IS was also analyzed on SAg-specific conjugates formed with CTLs where PLK1 expression was silenced by RNA interference (Fig.2B). PLK1 KD CTLs recapitulated the defect observed in BI2536-treated CTLs (Fig.2C), supporting the key role of PLK1 in IS formation. PLK1 blockade resulted in defective synaptic TCR accumulation also in Jurkat T cells and freshly purified peripheral blood CD4^+^ T cells (Fig.S2D,E).

**Figure 2.**
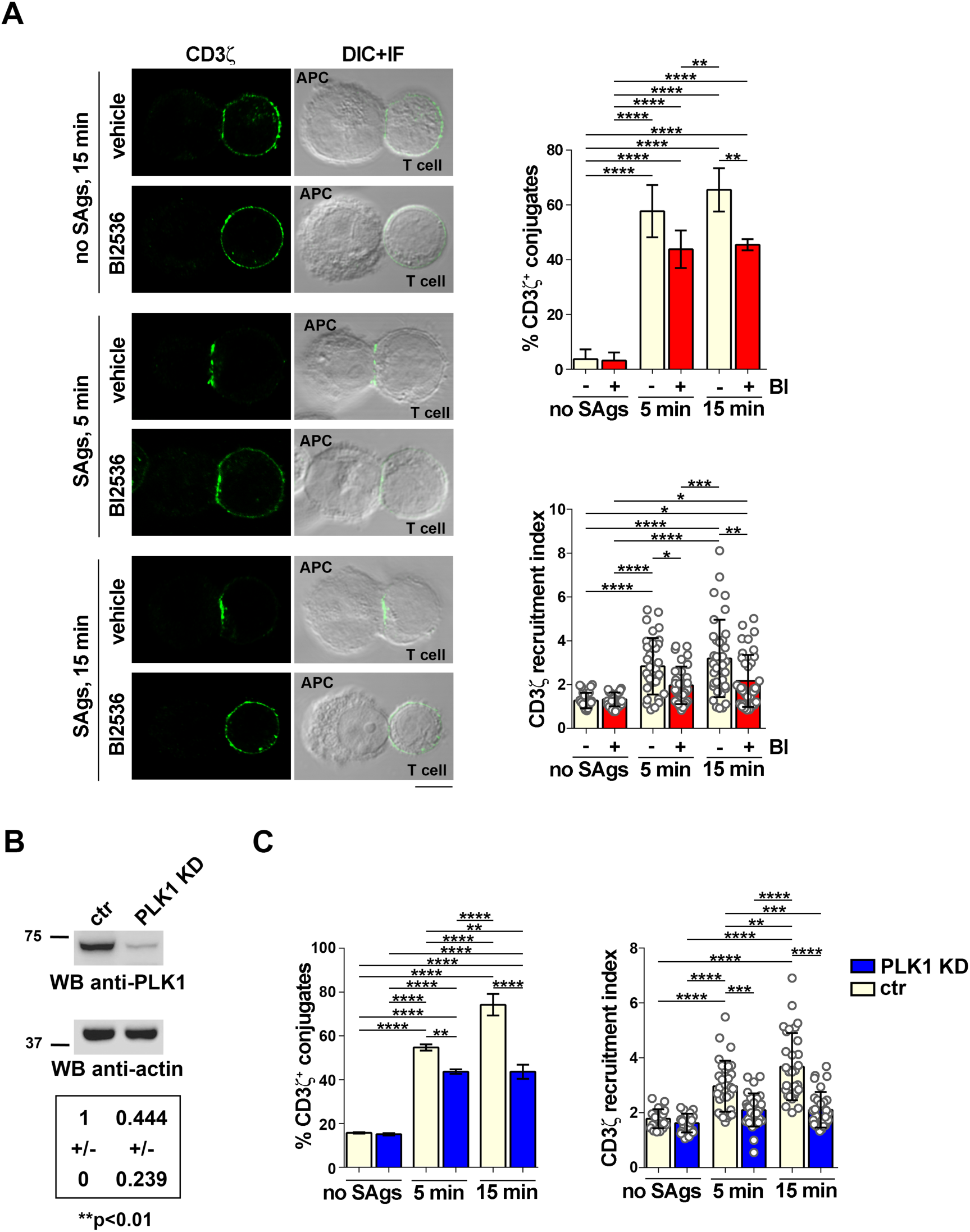
PLK1 inhibition or depletion impairs TCR clustering at the IS in CTLs. **(A)** Immunofluorescence analysis of the TCR subunit CD3ζ in 5 or 15 minutes conjugates of control (vehicle) and BI2536-treated CTLs and SAg-loaded Raji cells. Representative images of medial optical sections are shown. Scale bar: 5 μm. The histograms show the quantification of the percentages of conjugates positive for CD3ζ polarization to the IS (top) and the quantification of the relative CD3ζ fluorescence intensity at the IS (bottom)(20 cells/sample, n≥3; ANOVA). (**B**) Immunoblot analysis of PLK1 in representative matched control (scramble RNAi) and PLK1 KD CTLs, with respective loading control (actin) is shown. The migration of molecular mass markers is indicated. The quantification of PLK1 in CTL lysates is reported in the table (n=3; mean fold ± SD, one sample t test). (**C**) Quantification of the immunofluorescence analysis of 5 or 15 minutes conjugates of control and PLK1 KD CTLs and SAg-loaded Raji cells. The histograms show the percentages of conjugates positive for CD3ζ polarization to the IS (left) and the quantification of the relative CD3ζ fluorescence intensity at the IS (right) (10 cells/sample, n=3; ANOVA). *P < 0.05; **P < 0.01; ***P < 0.001; ****P < 0.0001.

The PLK1 upstream activator AurA has been implicated in early TCR signaling by promoting the activatory phosphorylation and synaptic localization of the initiating kinase Lck in CD4^+^ T cells (Blas-Rus et al., 2016). To understand whether PLK1 may mediate this effect of AurA in CTLs we imaged the active form of Lck, phosphorylated on Y394, in BI2536-treated CTLs conjugated with SAg-pulsed Raji cells. A defect in the IS accumulation of active Lck was observed in PLK1-inhibited CTLs (Fig.3A). Inhibition of Lck phosphorylation on Y394 in the presence of BI2536 was also observed by immunoblot analysis of post-nuclear supernatants from CTLs activated by antibody-mediated TCR cross-linking (Fig.3B). Consistent with this defect, immunoblot analysis of the active forms of two proteins that participate at sequential steps in the signaling cascade downstream of Lck, namely the transmembrane adaptor LAT and the kinase Erk1/2 (Samelson, 2002), showed that BI2536 treatment led to an impairment in TCR-dependent LAT and Erk phosphorylation (Fig.3C). The results indicate that PLK1 regulates the TCR signaling cascade in CTLs, accounting for its requirement for IS formation.

**Figure 3.**
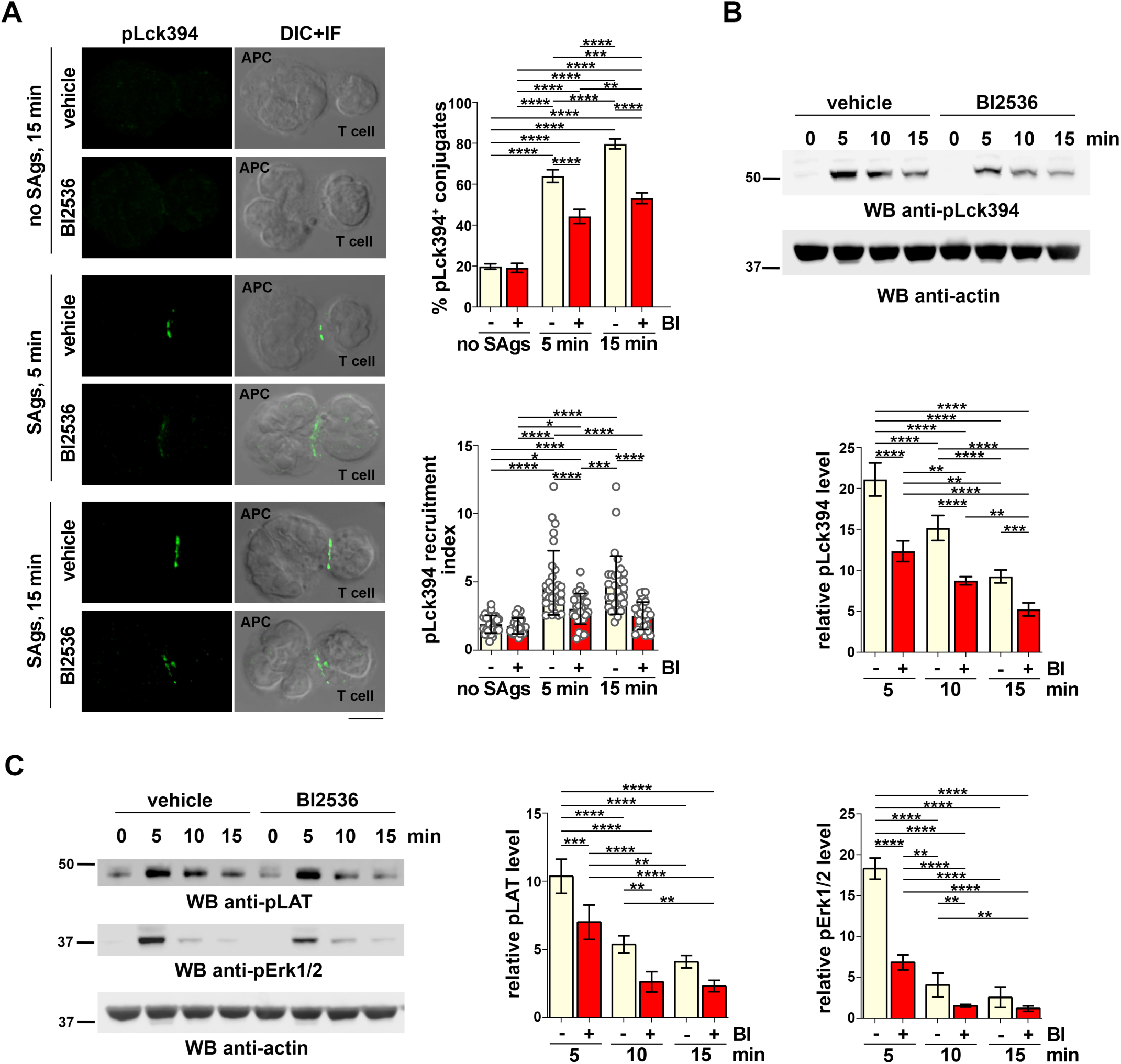
PLK1 inhibition impairs TCR signalling in CTLs. **(A)** Immunofluorescence analysis of the phosphorylated active form of Lck (Y394) (pLck394) in 5 or 15 minutes conjugates of control (vehicle) or BI2536-treated CTLs and SAg-loaded Raji cells. Representative images of medial optical sections are shown. Scale bar: 5 μm. The histograms show the quantification of the percentages of conjugates positive for pLck394 polarization to the IS (top) and the quantification of the relative pLck394 fluorescence intensity at the IS (bottom) (10 cells/sample, n≥3; ANOVA). **(B,C)** Immunoblot analysis with anti-pLck394 (B), anti-pLAT and anti-pErk1/2 (C) antibodies of lysates of either control or BI2536-treated CTLs, unstimulated or stimulated with anti-CD3 mAb for the indicated times. A control anti-actin blot of the stripped filter is shown below. The migration of molecular mass markers is indicated. The graphs show the quantification of pLck394 (B), pErk1/2 and pLAT (C), respectively (nβ3; mean fold ± SD, one-way ANOVA). *P < 0.05; **P < 0.01; ***P < 0.001; ****P < 0.0001.

### PLK1 is required for centrosome and LG polarization to the IS and for CTL-mediated killing

Centrosome translocation toward the T cell interface with cognate APC occurs in response to TCR engagement and in turn orchestrates the polarized recycling of endosome-associated TCRs to the synaptic membrane (Das et al., 2004). In CTLs, centrosome reorientation toward the IS, concomitant with the dynamic reorganization of the microtubule cytoskeleton, ensures the polarized transport of LGs along the microtubules to the site of contact and the release of their contents into the synaptic cleft (Douanne and Griffiths, 2021; Kabanova et al., 2018). Confocal analysis of SAg-specific conjugates formed with BI2536-treated CTLs showed an impairment in both centrosome and LG repositioning beneath the IS membrane, as assessed by quantifying both the frequency of conjugates harboring centrosome and LG polarization at the IS and their distance from the synaptic membrane (Fig.4A,B,C). Similar results were obtained using PLK1 KD CTLs (Fig.4E,F) and a CMV-specific CTL clone (Fig.S1D,E,F). Furthermore, LG convergence around the centrosome, which occurs in preparation for their transport to the IS (Calvo and Izquierdo, 2021), was impaired in BI2536-treated CTLs, as assessed by measuring the distance of individual GzmB^+^ vesicles from the PCNT-1 positive compartment (Fig.4D). This defect was recapitulated in PLK1 KD CTLs (Fig.4G). The inhibitory effect of BI2536 on centrosome polarization was also observed in CMV-specific CTL clone (Fig.S1G), Jurkat and freshly purified peripheral CD4^+^ T cells (Fig.S2D,E).

**Figure 4.**
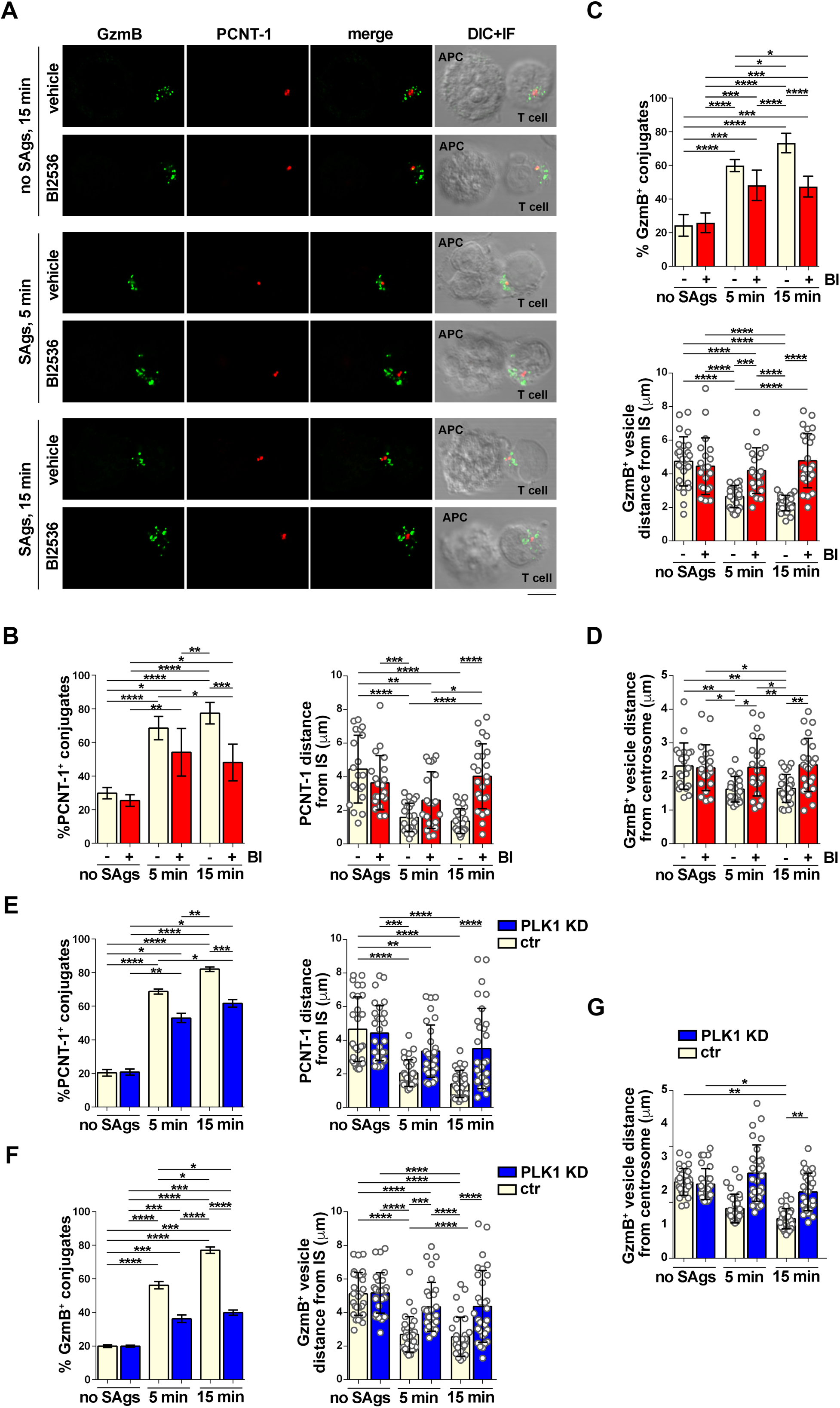
PLK1 inhibition impairs centrosome and LG polarization to the IS in CTLs. **(A)** Immunofluorescence analysis of the centrosomal protein pericentrin (PCNT-1) and the LG component granzyme B (GzmB) in 5 and 15 minutes conjugates of vehicle- and BI2536-treated CTLs and SAg-pulsed Raji cells. Medial optical sections of representative images are shown. Scale bar: 5 μm. **(B,E)** Quantification of the percentages of conjugates positive for PCNT-1 at the IS (left) and of the distance of the centrosome from the synaptic membrane (right) (μm) in conjugates of control or BI2536-treated CTLs and Raji B cells (B) or of control and PLK1 KD CTLs (E). **(C,F)** Quantification of the percentages of conjugates positive for GzmB at the IS (top (C), left (F)) and of the mean distance of single GzmB^+^ vesicles from the synaptic membrane (bottom (C), right (F)) (μm) in individual conjugates of control or BI2536-treated CTLs (C), or of control and PLK1 KD CTLs (F) and Raji B cells. **(D,G)** Quantification of the mean distance of single GzmB^+^ vesicles from the centrosome in individual conjugates of control and either BI2536-treated (D) or PLK1 KD (G) CTLs and Raji B cells (μm). Data are reported as mean ± SD. Statistical analysis: B-D: 20 cells/sample, n≥3; E-G: 10 cells/sample, n=3; ANOVA. *P < 0.05; **P < 0.01; ***P < 0.001; ****P < 0.0001.

### PLK1 promotes microtubule growth in CTLs

Upon TCR stimulation, microtubule filaments and microtubule motor proteins anchored at the cell cortex at the CTL interface with its cell target provide the pulling forces for centrosome translocation (Kuhn and Poenie, 2002). Microtubules are large polymers composed of α/β-tubulin dimers that are endowed with an intrinsic polarity, with (-) ends anchored to the centrosome to avoid rapid depolymerization and (+) ends characterized by dynamic instability (Li and Gundersen, 2008). After centrosome polarization, microtubules actively polymerize at the IS. Altering microtubule (+) end dynamics delays centrosome reorientation and vesicular traffic to the IS (Hooikaas et al., 2020; Martín-Cófreces et al., 2012), underscoring the importance of microtubule plasticity in the establishment of a functional IS.

PLK1 plays a crucial role in orchestrating microtubule dynamics in cycling cells (Zitouni et al., 2014). We hypothesized that the architecture of the microtubule cytoskeleton might be altered in PLK1-inhibited CTLs. To address this point, we imaged control and BI2536-treated CTLs plated under non-activating conditions (poly-L-lysine) and stained for α-tubulin. Control CTLs displayed numerous thin microtubules radiating from the centrosome (Fig.5A,B). When PLK1 activity was inhibited, microtubules organized in thick bundles that in some cases appeared to originate in the cortical area (Fig.5A,B). To quantify this feature we measured the skewness of pixel intensity, an amply validated method for the quantitative detection of bundling of actin filaments and microtubules (Higaki, 2010; Higaki, 2017). The skewness of the fluorescence intensity distribution becomes higher when the number of pixels with high fluorescence intensity is increased, providing a quantitative and reliable measurement of cytoskeleton bundling. BI2536-treated CTLs showed a higher skewness in α-tubulin staining compared to control cells (Fig.5C), providing evidence that PLK1 is involved in the regulation of the architecture of the microtubule cytoskeleton in CTLs.

**Figure 5.**
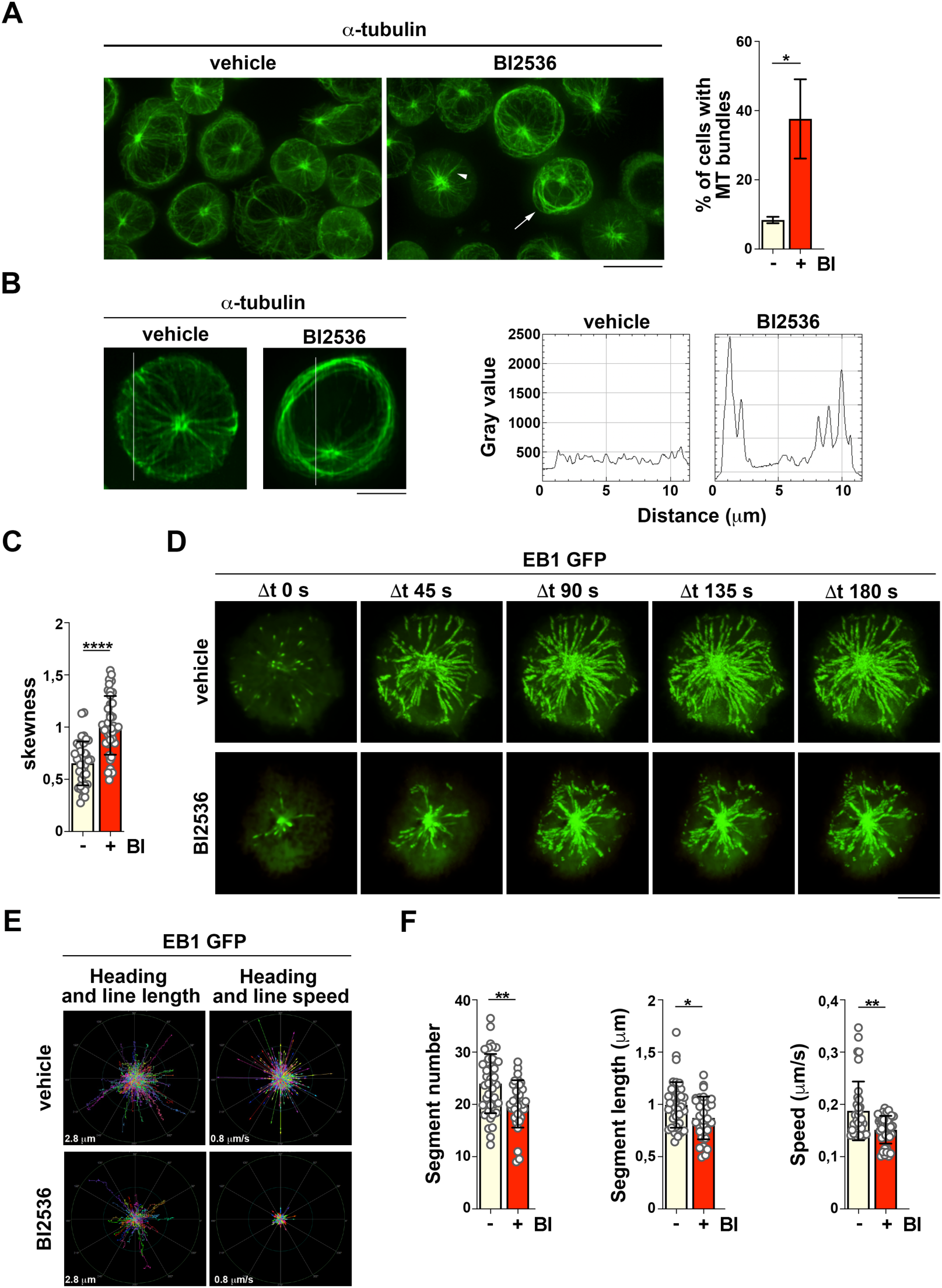
PLK1 inhibition impairs microtubule dynamics in CTLs. (**A,B**) Immunofluorescence analysis of α-tubulin in control and BI2536-treated CTLs. Representative images of maximum intensity projections of z-stacks are shown. Scale bar: 10 μm. (**A**) The arrowhead points to an example of cell displaying microtubules organized in few and thick α-tubulin-decorated bundles, while the arrow points to a cell showing microtubule bundles originating from the cortical area. The graph shows the percentage of cells displaying thick microtubule bundles (n=3; mean fold ± SD, t test). (**B**) Profiles of fluorescence intensity along a manually-selected line (white bar) in representative control or BI2536 treated CTLs. (**C**) Skewness of α-tubulin intensity distribution in control and BI2536-treated CTLs (240 cells from 3 independent experiments; mean fold ± SD, t test). (**D**) Imaging of EB1-EGFP-expressing CTLs, pre-treated with DMSO or BI2536 and plated on anti-CD3/CD28-coated glass-bottom chambers. Representative projections of time lapse are shown (βt = 45 sec). Images were taken every 300 ms for 3 minutes using a TIRF microscope. Scale bar: 5 μm. (**E**) Representative polar graphs of images in panel D showing the trajectories (left) and the speed (right) of EB1-EGFP-decorated microtubule (+) tips showing, respectively, the projection of object movements (color path length corresponds to track length) and the heading of each object as a color arrow (length corresponds to object speed). The external circle in the polar graphs indicates a distance of 2.8 μm (left) or a speed of 0.8 μm/s (right). (**F**) Quantification of the number (left) and length (center) of segments, and the speed of the growing tracks (right), reported as average value per cell (232 cells from 3 independent experiments; mean fold ± SD, t test). *P < 0.05; **P < 0.01; ***P < 0.001; ****P < 0.0001.

Since microtubule dynamics regulates the polarization of the centrosome to the IS (Stinchcombe and Griffiths, 2014), we asked whether the TCR signaling defect observed in BI2536-treated CTLs under steady-state conditions may impact on microtubule growth during IS assembly. CTLs were transiently transfected with a plasmid construct encoding EGFP-tagged EB1, a protein that binds to the (+) end of growing microtubules (Nehlig et al., 2017), and plated on glass-immobilized anti-CD3 mAb to induce centrosome polarization and microtubule polymerization (Bunnell et al., 2002). The dynamics of microtubule growth following CTL stimulation was tracked by EB1-EGFP live cell imaging and total internal reflection fluorescence (TIRF) microscopy to improve spatial and time resolution (Dixit and Ross, 2010) (Fig.5D). BI2536-treated EB1-EGFP transfected CTLs displayed altered microtubule dynamics (Fig.5D-F; Movies 1,2). In particular, the number and length of each segment, identified as the portion of each EB1-EGFP labelled track with a constant trajectory, were decreased in CTLs following BI2536 treatment, showing that the directionality of newly formed tracks was affected when PLK1 was inhibited (Fig.5D-F). Furthermore, the growth of microtubules was slower in BI2536-treated CTLs compared to control cells (Fig.5D-F). Taken together, these results indicate that PLK1 regulates microtubule dynamics during IS assembly in CTLs.

### PLK1 is required for efficient CTL-mediated killing

Rapid LG secretion upon target cell encounter can occur in the absence of centrosome translocation towards the IS membrane (Bertrand et al., 2013). Yet, the IS polarization of the CTL lytic machinery ensures the prolonged and confined secretion of lytic molecules onto the target cell, as demonstrated by the reduction in cytotoxicity when centrosome reorientation is delayed or defective (De La Roche et al., 2013; Jenkins et al., 2014; Tsun et al., 2011). To test the outcome of the IS defects associated with PLK1 inhibition on the killing capability of CTLs we performed a time course analysis of fluorescent calcein release by SAg-pulsed target cells induced by either BI2536-treated or untreated CTLs. In this assay target cells are loaded with the cell-permeant dye calcein AM, that becomes fluorescent following hydrolyzation by intracellular esterases and is released upon cell death (Chang et al., 2018). Consistent with the role of PLK1 in TCR signaling and IS assembly, the killing capability of CTLs was impaired at every effector:target cell ratio (E:T) tested when PLK1 activity was inhibited (Fig.6A). Similar results were obtained using PLK1 KD CTLs (Fig.6B). Hence PLK1 contributes to CTL-mediated killing through its ability to regulate IS assembly.

**Figure 6.**
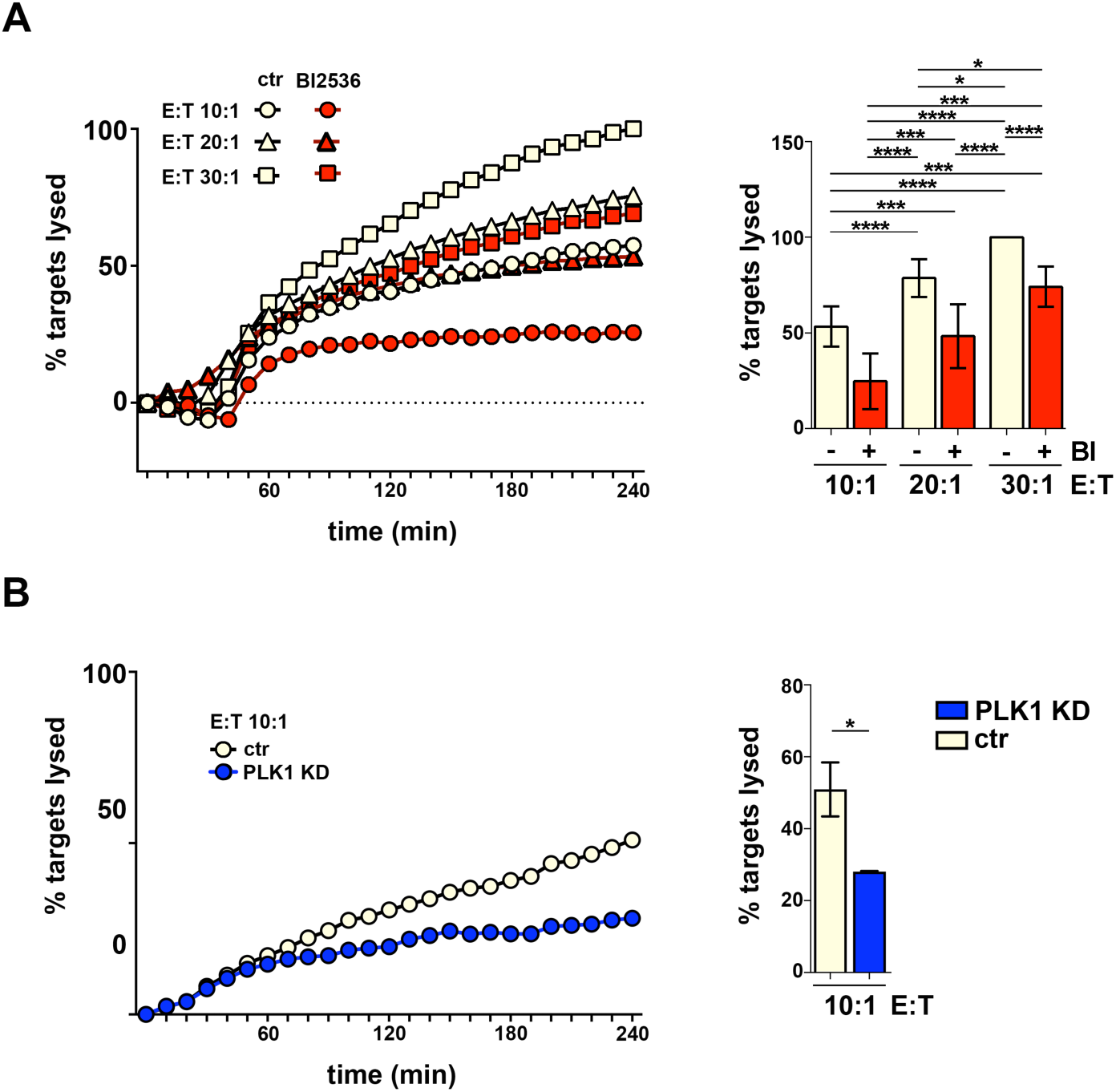
PLK1 inhibition impairs CTL-mediated killing. **(A,B)** Real-time calcein release-based killing assay. CTLs either untreated or pre-treated with BI2536 (A) and control or PLK1 KD CTLs (B), were co-cultured with SAg-loaded Raji B cells at the target:CTL ratios indicated, and target cell killing was measured every 10 minutes for 4 hours as reported in the kinetic graph (left). Quantification of the percentage of target cell death at the endpoint of the procedure (4 h) is shown on the right graph. The data refer to at least three independent experiments performed in duplicate and are reported as mean fold ± SD, with target cell lysis at the highest target:CTL ratio of the control sample set at 100% of cytotoxicity (ANOVA). *P < 0.05; **P < 0.01; ***P < 0.001; ****P < 0.0001.

## Discussion

The role of mitotic kinases in non-cycling cells has been as yet poorly investigated. Here we show that PLK1, known to regulate centrosome duplication and microtubule dynamics during mitosis (Joukov and De Nicolo, 2018), is required for IS assembly in non-cycling T cells. We provide evidence that PLK1 subserves a dual role in this process, participating in the TCR signaling cascade and modulating microtubule dynamics. Both these functions converge on centrosome polarization towards the T cell contact with its cognate target, a key step in the formation of a functional IS. In CTLs, the failure of the centrosome to polarize to the IS following pharmacological inhibition or silencing of PLK1 led to impaired LG translocation towards the target cell and defective target cell killing, identifying PLK1 as a new regulator of CTL function.

Key players in early TCR signaling, including the kinases Lck and ZAP-70 and the adaptors LAT and SLP-76, are required for centrosome polarization (Kuhné et al., 2003; Lowin-Kropf et al., 1998). How these early events translate into the physical movement of the centrosome towards the subsynaptic region has been in part elucidated. Diacylglycerol locally produced at the IS by PLCγ acts as a polarity cue for the centrosome through the recruitment of members of the novel and atypical PKC subfamilies (Huse, 2012). TCR signaling also promotes the synaptic recruitment of the dynein-dynactin complex that has been proposed to provide pulling forces on the microtubules radiating from the centrosome (Liu et al., 2013). Additionally, actin filaments that polymerize at the IS periphery interact with these microtubules to provide tension for centrosome translocation (Calvo and Izquierdo, 2021). The mechanism linking the novel and atypical PKCs to the cytoskeletal elements that provide the pulling forces remains to be fully characterized. The implication of PLK1 in TCR signaling, starting with the activation of the initiating kinase Lck, is likely to account, at least in part, for the defect in centrosome polarization observed in PLK1-inhibited or -depleted T cells. The TCR-dependent recruitment of a pool of phosphorylated PLK1 to the IS places it in a strategic position to regulate the early signaling events that orchestrate centrosome polarization.

How PLK1 regulates the activity of Lck remains to be determined. The activity of Lck is tightly controlled by the phosphorylation status of Y394 and Y505. While full activation requires Y505 dephosphorylation by the transmembrane tyrosine phosphatase CD45 and Y394 autophosphorylation, serine phosphorylation contributes to fine-tuning its activity (Rossy et al., 2012). The threonine/serine kinase Erk has been shown to phosphorylate Lck on S59 under conditions of strong TCR stimulation, which prevents the recruitment of the phosphatase SHP-1 that disables Lck by dephosphorylating Y394, thereby sustaining TCR signaling (Štefanová et al., 2003). PLK1 could be hypothesized to participate directly or indirectly in such regulatory circuitry. As the search for PLK1 substrates has been mainly restricted to the context of mitosis, extending these studies to non-cycling cells may provide new insights into the non-mitotic functions of this kinase.

In mitotic cells PLK1 regulates microtubule dynamics by promoting microtubule nucleation at the centrosome through phosphorylation of PCM components, such as SPD-5 in *C. elegans*, that recruit the γ-tubulin ring complex (γ-TuRC) (Ohta et al., 2021). Additionally, PLK1 regulates microtubule-based microtubule nucleation in human cells during mitotic spindle formation by promoting the interaction of the γ-TuRC recruitment protein, Nedd1, with microtubule-bound Augmin (Johmura et al., 2011). PLK1 has also been implicated in the control of microtubule stability in *D. melanogaster* and mammalian cells through its ability to modulate the activity of the microtubule-associated protein CLASP and the (+) end microtubule motor Kif13 (Moriwaki and Goshima, 2016). Consistent with a role for PLK1 in microtubule dynamics, the rate and extent of microtubule growth has been shown to decrease in PLK1-inhibited human breast adenocarcinoma MCF-7 cells, concomitant with an increase in microtubule acetylation (Rashid et al., 2020). Microtubule dynamics is essential for centrosome polarization to the IS. Genetic depletion of regulators of microtubule growth or stability in T cells, such as the microtubule-binding proteins EB1 and stathmin (Filbert et al., 2012; Martín-Cófreces et al., 2012; Zyss et al., 2011) or modulation of tubulin acetylation through HDAC6 overexpression (Serrador et al., 2004), have been associated with impairment or delay in centrosome polarization to the IS and defective T cell function. It is noteworthy that, while faster rates of microtubule growth translate to increased traction forces that improve the killing activity of CTLs (Pathni et al., 2022), microtubule overgrowth, as observed in T cells depleted of the kinesin Kif21B, also leads to a similar defect (Hooikaas et al., 2020), underscoring the notion that microtubule dynamics must be tightly controlled during the complex process of centrosome translocation to the IS and the subsequent polymerization of new microtubules from the repositioned centrosome. Our finding that PLK1 regulates microtubule growth during IS assembly suggests that its ability to promote centrosome polarization might not be limited to its role in TCR signaling but also depend on a more direct role in microtubule nucleation at the centrosome. Interestingly, the *Xenopus* Polo-like kinase Plx1 has been reported to regulate the function of stathmin/Op18 (Budde et al., 2001). Additionally, in pre-mitotic ciliated cells PLK1 promotes cilium disassembly by phosphorylating HDAC6, thereby promoting microtubule deacetylation and disassembly (Wang et al., 2013). The finding that EB1, stathmin and HDAC6 are recruited to the IS in T cells (Filbert et al., 2012; Martín-Cófreces et al., 2012; Serrador et al., 2004) provides a potential direct link between PLK1 and microtubule dynamics during IS assembly.

It is noteworthy that the effects of PLK1 inhibition or depletion in T cells reported here do not completely phenocopy the effects elicited by blockade or depletion of its upstream regulator AurA in CD4^+^ T cells (Blas-Rus et al., 2016). Both associate with the T cell centrosome and localize at the IS, where a pool of either active kinase can be found. They are both required for Lck activation and early TCR signaling and for microtubule growth during IS formation. However, a major difference is in their ability to promote centrosome translocation, which is not affected in AurA-targeted T cells (Blas-Rus et al., 2016). While the study on AurA was carried out on freshly purified CD4^+^ T cells and our study mainly on CD8^+^ cells differentiated to CTLs, we found a similar inhibitory effect of PLK1 blockade on centrosome polarization to the IS in freshly purified CD4^+^ T cells, as well as in the CD4^+^ T cell-derived Jurkat cell line. This suggests that, while the AurA/PLK1 axis that regulates mitosis is operational during IS assembly in non-cycling T cells, PLK1 does not only act as effector of AurA but also subserves AurA-independent functions in this process.

Not surprisingly considering its key role in cell division, PLK1 is overexpressed in many types of both solid and blood cancers (Iliaki et al., 2021). A plethora of studies have provided evidence that targeting PLK1 using selective pharmacological inhibitors blocks the proliferation of cancer cells and promotes their apoptosis, setting the basis for a number of clinical trials (Gutteridge et al., 2016; Park et al., 2015; Strebhardt, 2010; Su et al., 2022; Zeidan et al., 2020). Our finding that PLK1 is required for IS assembly, on which T cell activation depends, highlights a potential drawback of PLK1-targeted therapies. The T cell suppressive activity of a PLK1-specific inhibitor on the activation of alloreactive T cells in acute graft-versus-host disease (Hooikaas et al., 2020), while highlighting the potential therapeutic benefits of targeting PLK1 in this disease context, underscores the importance of investigating the immune functions of cancer patients undergoing PLK1-directed therapies.

## Materials and methods

### Cells, T cell transfectants, antibodies

CD8^+^ and CD4^+^ T cells were isolated from peripheral blood of anonymous healthy donors obtained from the Siena University Hospital blood bank. The study was approved by the local ethics committee (Siena University Hospital). Informed consent was obtained from blood donors by the physician in charge of the Siena University Hospital blood bank. Samples were anonymized before distribution. Primary CD8^+^ and CD4^+^ T cells were isolated by negative selection through either RosetteSep^TM^ Human CD8^+^ T Cell Enrichment Cocktail (StemCell technologies, Vancouver, Canada) or RosetteSep^TM^ Human CD4^+^ T Cell Enrichment Cocktail (StemCell technologies, Vancouver, Canada) and centrifugation over a buoyant density medium (Lympholyte Cell Separation Medium, Euroclone, Milan, Italy). Immediately after isolation, CD8^+^ and CD4^+^ T cells were resuspended in RPMI 1640 with 25 mM Hepes (Sigma-Aldrich, St. Louis, Missouri, USA), supplemented with 10% BCS (Bovine Calf Serum, Hyclone, Logan, Utah, USA) inactivated at 56 °C for 30 minutes, 20 U/mL Penicillin (Sigma-Adrich, St. Louis, Missouri, USA) and 1X non-essential amino acids (MEM non-essential amino acids solution 100X, Gibco, Waltham, Massachusetts, USA). Cell cultures were maintained at 37 °C and 5% CO_2_. Freshly isolated CD8^+^ T cells were differentiated *in vitro* to CTLs by incubation for 48 h with anti-CD3/CD28 coated magnetic beads (Dynabeads Human T-activator CD3/CD28, Gibco, Waltham, Massachusetts, USA) for cell expansion and activation and 50 U/mL of IL-2 (human Interleukin-2, Miltenyi Biotech, Bergisch Gladbach, Germany) for cell proliferation and differentiation. Mature CTLs (days 5 to 7 of differentiation) were used for the experiments (Onnis *et al*., 2022). Freshly isolated CD4^+^ T cells were immediately used for the assays.

To generate CTL clones, CD8^+^ T cells specific for the VLAELVKQI peptide of the cytomegalovirus protein pp65 were single cell sorted into 96-U-bottom plates using a BD FACSAria II cell sorter using tetramer staining. Cells were cultured in RPMI 1640 medium supplemented with 8% human AB serum (PAA), minimum essential amino acids, HEPES and sodium pyruvate (Invitrogen), 100 IU/ml human rIL-2 and 50 ng/ml human rIL-15. CD8^+^ T-cell clones were stimulated in complete RPMI/HS medium containing 1 mg/ml PHA with 1 x 10^6^/ ml 35 Gy irradiated allogeneic peripheral blood mononuclear cells (isolated on Ficoll Paque Gradient from buffy coats of healthy donors) and 1 x 10^5^/ ml 70 Gy irradiated EBV-transformed B cells. Re-stimulation of clones was performed every 2 weeks. Blood samples were collected and processed following standard ethical procedures (Helsinki protocol), after obtaining written informed consent from each donor and approval for this study by the local ethical committee (Comité de Protection des Personnes Sud-Ouest et Outremer II). Other cells used were Raji B cells, EBV-transformed B cells (JY) and Jurkat T cells (ATCC, Manassas, VA), maintained in RPMI 1640 supplemented with 7.5% BCS and 20 U/ml Penicillin.

For RNAi-mediated PLK1 silencing CTLs were transiently transfected at day 6 of differentiation using the Human T cell nucleofector kit and the program T-023 of the Nucleofector II system (Amaxa Biosystems, Euroclone, Milan, Italy) for activated cells with human PLK1-specific siRNAs (#4390824, s448) and negative control siRNA (#4390846) (Invitrogen, Waltham, Massachusetts, USA) (150 ng/10^6^ cells). Cells were used 24 h post-transfection. All samples were tested by immunoblotting to check the efficiency of PLK1 knockdown.

CTLs were transiently transfected with the pEGFP plasmid encoding the EB1-GFP fusion protein (Addgene #17234, Watertown, Massachusetts, USA). Briefly, CTLs at 5 days of differentiation were transiently transfected using the Human T cell nucleofector kit and the program T-023 of the Nucleofector II system (Amaxa Biosystems, Euroclone, Milan, Italy) for activated cells with 1 μg/10^6^ cells of pEGFP-EB1 plasmid. Transfected cells were gently resuspended in the culture medium added with 500 U/ml IL-2 and analysed 24 h post-transfection.

T cells were pre-treated with the PLK inhibitor BI2536 (Selleckchem, Houston, Texas, USA) at the concentration of 100 nM in DMSO in serum-free medium for 3 h at 37°C and 5% CO_2_ and used as such for the assays (Lénárt et al., 2007). Control samples were incubated with the same amount of DMSO.

All primary commercial antibodies used in the assays are listed in Table 1, together with information on the dilutions used for immunoblotting and immunofluorescence. Secondary horseradish peroxidase (HRP)-labelled antibodies were purchased from Jackson ImmunoResearch Laboratories (West Grove, Pennsylvania, USA) and Alexa Fluor 488- and 555-labeled secondary antibodies from ThermoFisher Scientific (Waltham, Massachusetts, USA). IgG from OKT3 (anti-human CD3ɛ, IgG2) hybridoma supernatants was purified using Mabtrap (Amersham Biosciences, Inc., Piscataway, NJ, USA) and titrated by flow cytometry.

### Conjugate formation, CTL activation on activating surfaces, immunofluorescence and image analysis

Conjugates between CD8^+^ T cells and superantigen-pulsed Raji B cells were carried out as previously described (Cassioli et al., 2021). Raji B cells (used as APCs) were loaded with 10 μg/ml of a mix of Staphylococcal Enterotoxins A (SEA), B (SEB) and E (SEE) (Toxin Technologies, Sarasota, Florida, USA) for 2 h at 37°C and labelled with 10 μM Cell Tracker Blue for the last 20 min of the incubation with superantigens (SAgs). Conjugates of CTLs with unpulsed Raji B cells were used as negative controls. SAg-pulsed or unpulsed Raji B cells were mixed with CTLs (1:1) to allow conjugate formation at 37°C for the indicated time points. Samples were seeded onto poly-L-lysine (Merck, Drmstadt, Germany)-coated slides (ThermoFisher Scientific, Waltham, Massachusetts, USA). Alternatively, CTLs were activated in the absence of target cells by plating on 5 μg/ml anti-CD3 antibody (clone OKT3)(Bio Legend, San Diego, California, USA)-coated slides. Cells were fixed with either methanol at -20°C for 10 min or 4% paraformaldehyde/PBS at room temperature for 15 min. After washing with PBS, cells were stained with primary antibodies at 4°C overnight, and then incubated at room temperature for 45 min with Alexa fluor 488- and 555-labeled secondary antibodies and mounted with 90% glycerol/PBS.

Confocal microscopy was carried out on a Zeiss LSM700 microscope (Carl Zeiss, Jena, Germany) using a 63x/1.40 oil immersion objective or a spinning disk confocal and super-resolution microscope (CSU-W1-SoRA Nikon), with 100x/1.49 oil objective. 3D deconvolution (Blind method, 20 iterations) and denoise were performed using the software NIS Elements AR Nikon and applied to high-resolution images. Detectors were set to detect the optimal signal below the saturation limits.

Co-localization analyses were performed on medial optical sections of single cells using ImageJ and the JACoP plugin to calculate Mander’s coefficient (Manders et al., 1992). Mander’s coefficients range from 0 to 1, corresponding to non-overlapping images and 100% co-localization between both images, respectively.

Polarization of CD3ζ, GzmB^+^ lytic granules and PLK1, and the translocation of the centrosome to the IS, were based on the presence of the staining solely on the CTL:APC contact site and were expressed as the percentage of conjugates with synaptic staining *versus* the total number of conjugates analysed. The relative distance of the centrosome, PLK1, AurA and individual GzmB^+^ vesicles from the contact site of CTLs with APCs or individual GzmB^+^ vesicles from the centrosome was measured using ImageJ. The recruitment index was calculated as the ratio of either CD3ζ or pPLK1 fluorescence intensity at the synaptic area, which is manually defined at the CTL:APC contact site *versus* the entire cell membrane, using ImageJ. The relative pPLK1 fluorescence at the centrosome was calculated over a circular region defined from the point of PCM1 maximal intensity. The skewness of the intensity distribution of α-tubulin was measured using Fiji, by calculating the mean value of 3 different regions (2×3 μm) for each cell, excluding the centrosomal area.

### Live cell imaging and analysis

For TIRF microscopy, EB1-EGFP transfected CTLs were washed and seeded at 1.5 x 10^5^ cells per well on 5 μg/ml anti-CD3 antibody (clone OKT3)-coated eight-well chambered slides (Ibidi, Munich, Germany). Chambered slides were mounted on a heated stage within a temperature-controlled chamber maintained at 37 °C, and constant CO_2_ concentrations (5%). Cells were visualized using a CSU-W1-SoRA Nikon microscope coupled to Photometrics BSI (Nikon) camera fitted with 100x/1.49 oil objective. Images were taken every 300 ms for 3 minutes and time lapses were processed with the confocal software NIS Elements AR Nikon. Images were equalized in intensity in time, denoised, background corrected and then bright spots were detected and tracked.

### Cell lysis and Immunoblotting

Cells (2×10^6^/sample) were stimulated with purified anti-human CD3ɛ for the indicated time at 37°. Cells were then lysed in 0.5% (v/v) Triton X-100 in 20 mM Tris–HCl (pH 8), 150 mM NaCl in the presence of Protease inhibitor Cocktail Set III (Calbiochem®, Merck, Darmstadt, Germany) and the phosphatase inhibitor sodium orthovanadate (Sigma-Aldrich, St. Louis, Missouri, USA) for 5 min on ice. Protein extracts from post-nuclear supernatants were quantified with the Quantum protein assay kit (Euroclone, Milan, Italy) and denatured in 4x Bolt SDS sample buffer (Invitrogen, Waltham, Massachusetts, USA) supplemented with 10x Bolt sample reducing buffer (Invitrogen, Waltham, Massachusetts, USA) for 5 min at 100°C. Proteins (10 μg) were subjected to SDS-PAGE on Bolt Bis-Tris mini protein gels (Invitrogen, Waltham, Massachusetts, USA) and transferred to nitrocellulose (GE HealthCare, Euroclone, Milan, Italy) under wet conditions. Blocking was performed in 5% non-fat dry milk in PBS containing 0.2% Tween 20 (Sigma-Aldrich, St. Louis, Missouri, USA). Membranes were incubated in primary antibodies for 1-3 h at room temperature (20-25°C) or overnight at 4°C, followed by incubation with 20 ng/ml HRP-conjugated secondary antibodies (Jackson ImmunoResearch Laboratories, West Grove, Pennsylvania, USA) for 45 min at room temperature. Secondary antibodies were detected using SuperSignal west pico plus chemiluminescent substrate (Life Technologies, Waltham, Massachusetts, USA). For quantification, immunoblot membranes were scanned using Alliance Q9-Atom chemiluminescence imaging system (Uvitec, Cambridge, UK), and densitometric levels were measured using ImageJ software (National Institutes of Health, USA).

### Real-time calcein release-based killing assay

Raji B cells were loaded with 500 nM calcein-AM (Life Technology, Waltham, Massachusetts, USA) in AIM V medium (ThermoFisher Scientific, Waltham, Massachusetts, USA) with 10 mM Hepes at room temperature for 15 min, washed and plated in 96-well black plates with clear bottom (BD Falcon, Corning, New York, USA). BI2536-treated or PLK1 KD CTLs and controls were added at different ratios to 0.5×10^4^ settled target cells per well to measure killing at 37°C, 5% CO_2_. Triton X-100 (1%) was added to target cells alone to calculate maximal target cell lysis as control. Target cell lysis was measured every 10 min for 4 h. The decreased calcein fluorescence in target cells due to cell lysis was measured at 485 nm excitation wavelength and 528 nm emission wavelength in the bottom reading mode using a Synergy HTX multi-mode plate reader (BioTek, Santa Clara, California, USA). The fluorescence for the experimental condition was adjusted by the parameter γ according to the live target cell control fluorescence. The γ value was measured at time zero: γ = F_live_(0)/F_exp_(0). Cytotoxicity was calculated based on the loss of calcein fluorescence in target cells using the equation: % target cell lysis = (F_live_ - γ × F_exp_)/(F_live_-F_lyse_) × 100, where F_live_ is the fluorescence of target cells alone, F_exp_ are CTL+APC samples and F_lyse_ is the maximal target cell lysis. All the experiments were performed in duplicate and averaged to obtain one dataset. The maximal CTL-induced target cell killing was assigned to the higher CTL:APC ratio, on which the relative values for the other samples were based (Chang et al., 2018).

### Statistics and reproducibility

Each experiment is the result of at least 3 independent replicates. The number of cells analysed is specified in the figure legends. Statistical analyses were performed using Prism software (GraphPad Software). Pairwise or multiple comparisons of values with normal distribution were carried out using Student’s t-test (unpaired), one-sample t-test (theoretical mean=1) and one-way ANOVA, whereas values without Gaussian distribution were analysed with Mann-Whitney test or Kruskal-Wallis test. Statistical significance was defined as: ****P≤0.0001; ***P≤0.001; **P≤0.01; *P≤0.05; n.s., not significant.

**Table T1.** List of the primary antibodies used in this work.

| Antibody (anti-) | Host species | Company | Catalog number | Dilution WB | Dilution IF |
| --- | --- | --- | --- | --- | --- |
| Actin | mouse | Millipore (Burlington, Massachusetts, USA) | MAB1501 | 1:10,000 | - |
| Aurora-A | rabbit | Cell signalling (Danvers, Massachusetts, USA) | 14475 | - | 1:100 |
| CD3ζ | mouse | Santa Cruz (California, USA) | sc-1239 | - | 1:200 |
| pErk1/2 (T202/Y204) | rabbit | Cell signalling (Danvers, Massachusetts, USA) | 9101 | 1:1,000 | - |
| Granzyme B-488 | mouse | Bio Legend (San Diego, California, USA) | 560211 | - | 1:20 |
| pLat (Y191) | rabbit | Cell signalling (Danvers, Massachusetts, USA) | 3584S | 1:1,000 | - |
| Lck | mouse | BD (Franklin Lakes, New Jersey, USA) | 610097 | - | 1:50 |
| pLck (Y394) | mouse | Bio-technie (Minneapolis, Minnesota, USA) | MAB7500 | 1:1000 | 1:50 |
| PCM1 | mouse | Santa Cruz<br>(California,<br>USA) | sc-398365 | - | 1:300 |
| Pericentrin | rabbit | Abcam<br>(Cambridge,<br>UK) | ab4448 | - | 1:200 |
| PLK1 | rabbit | Bio-technie<br>(Minneapolis,<br>Minnesota,<br>USA) | 104302 | 1:500 | 1:300 |
| pPLK1 (S137) | rabbit | Abcam<br>(Cambridge,<br>UK) | ab21738 | 1:1,000 | 1:300 |
| α-tubulin | mouse | Santa Cruz<br>(California,<br>USA) | sc-32293 | - | 1:500 |
| γ-tubulin | mouse | Sigma-Aldrich<br>(St. Louis,<br>Missouri, USA) | T6557 | - | 1:200 |

## Acknowledgements

The authors wish to thank Giuliano Callaini for critical reading of the manuscript and Annamaria Fusillo for technical assistance.

## Competing interests

The authors declare no competing financial interests.

## Funding

This research has received funding from the European Commission (ERC_2021_SyG 951329 - ATTACK) to CTB and SV and AIRC (IG 2017-20148) to CTB.

## Data availability

Data generated in this study are available upon request from corresponding author FF.

## Notes

### Competing Interest Statement

The authors have declared no competing interest.

